# Microbiopsy of living mouse brain for longitudinal molecular profiling

**DOI:** 10.64898/2026.01.22.701044

**Authors:** Alexander Hoyningen, Anna Ramisch, Loélia Fellouse, Agnès Hiver, Alma Lingenberg, Christian Lüscher, Lucile Marion-Poll

## Abstract

Longitudinal molecular studies of the mouse brain are limited by the need for terminal tissue collection. This prevents analysis of preexisting molecular states and their evolution within the same individual. We developed a stereotactic microbiopsy technique that enables minimally invasive sampling of defined brain regions *in vivo*. The method preserves survival while yielding material suitable for RNA and nuclei isolation. It provides a practical solution for linking baseline molecular states to subsequent behavioural, pharmacological, or disease-related outcomes.

**SUMMARY:** This study presents a stereotactic microbiopsy technique for sampling defined brain regions in living mice, enabling transcriptomic and epigenomic analyses without sacrificing the animal. The method will allow pre-intervention tissue collection, making it possible to separate preexisting molecular differences from experience- or treatment-induced changes. We show that microbiopsies yield sufficient, high-quality RNA and chromatin for sequencing, with minimal tissue damage that largely resolves over time. The procedure uses standard stereotactic equipment and achieves reproducible spatial precision when the syringe is stabilised. This approach provides a practical framework for within-subject molecular comparisons, reducing animal use and enabling longitudinal profiling of the living mouse brain. It establishes a foundation for investigating how baseline molecular states influence later physiological or behavioural outcomes.

## INTRODUCTION

Individual brain identity and function emerge from the interplay between genetic sequence, developmental stochasticity, experience-driven circuit activity and environmental influences. Activity-dependent transcription and epigenomic regulation are central to long-term circuit plasticity and are thought to contribute to inter-individual variability, even among genetically identical animals^1-3^. Mice represent the most widely used experimental model in neuroscience; however, molecular profiling in this species still relies almost exclusively on terminal tissue collection. This provides only a single molecular snapshot per animal and precludes the determination of whether observed differences reflect preexisting molecular configurations or arise as a consequence of experience.

The study of gene expression and chromatin accessibility in the brain have been revolutionised by high-throughput sequencing methods such as RNA sequencing (RNA-seq) and Assay for Transposase-Accessible Chromatin sequencing^4^ (ATAC-seq). Importantly, methodological developments now allow these analyses to be performed from very limited material, including low-input and even single-cell samples^5-7^. Nevertheless, despite their increased sensitivity, these approaches still rely on brain postmortem sampling and therefore cannot capture the dynamics of molecular states within the same individual over time.

Stereotactic brain biopsy represents an opportunity for brain pre-sampling. It is a well-established clinical procedure in humans (notably for tumors and encephalitis)^8^ and has also been adapted to large-animal models such as swine, allowing accurate sampling of deep brain structures with minimal risk^9^. Standard human biopsy needles typically retrieve 10–100 mg of tissue per core, whereas the entire mouse brain weighs approximately 400 mg. Thus, such biopsies cannot be used in mice as they would remove a large proportion of total brain mass. The degree of miniaturisation required to achieve tissue preservation and postoperative survival in mice consequently demands the development of different sampling tools.

In rodents, several *in vivo* sampling techniques have been employed, including microdialysis^10^, solid-phase microextraction^11^, and open-flow microperfusion^12^. While these methods permit the measurement of extracellular metabolites or drugs, they do not recover intact cells and are therefore unsuitable for nucleic acid–based assays. Biopsies have been described in tumor-bearing animals^13,14^, but these applications remove large volumes of pathological tissue and cannot be applied to healthy deep-brain regions. Consequently, to the best of our knowledge, there is currently no established method enabling recovery of high-quality RNA or nuclei from the healthy living mouse brain while maintaining anatomical integrity and animal survival.

Here, we describe a stereotactic microbiopsy technique that overcomes these limitations by allowing minimally invasive sampling of spatially defined brain regions *in vivo*. Using a blunt needle connected to a syringe, nanoliter-scale tissue volumes are aspirated under anaesthesia and frozen or immediately processed for RNA or nuclei isolation. We systematically quantify aspirated volume across needle gauges, evaluate tissue recovery over time, and assess targeting precision using atlas-based coordinates^15^. The material obtained yields high-quality RNA and intact nuclei, compatible with RNA-seq and ATAC-seq. This approach extends standard stereotactic surgery to molecular sampling and establishes a framework for longitudinal molecular analysis within individual mice, offering a means to relate baseline molecular states to subsequent functional or behavioural outcomes while simultaneously reducing animal use.

## RESULTS

### Design and implementation of stereotactic microbiopsy in mice

We developed a stereotactic microbiopsy method compatible with standard neurosurgical equipment (Fig. 1A–C), to enable molecular sampling from the living mouse brain (Fig. 1D). Animals were anesthetised and head-fixed in a standard stereotactic frame. A sterile blunt-tipped needle connected to a syringe was positioned at bregma to zero the stereotaxic coordinates. The needle was then moved above the target coordinate and a burr hole was drilled to access the cortical surface (Fig. 1E). During descent, positive pressure is maintained to prevent tissue located above the target from entering the needle lumen. Once the tip reaches the desired coordinate, it is left in place for ∼5 min until the entry site appears sealed and dry (Fig. 1F,G), supporting local vacuum formation during aspiration. Negative pressure is then applied by retracting the syringe plunger while manually stabilising the syringe to minimise needle deflection (Fig. 1H), aspirating a small, spatially defined tissue fragment.

**Figure 1:**
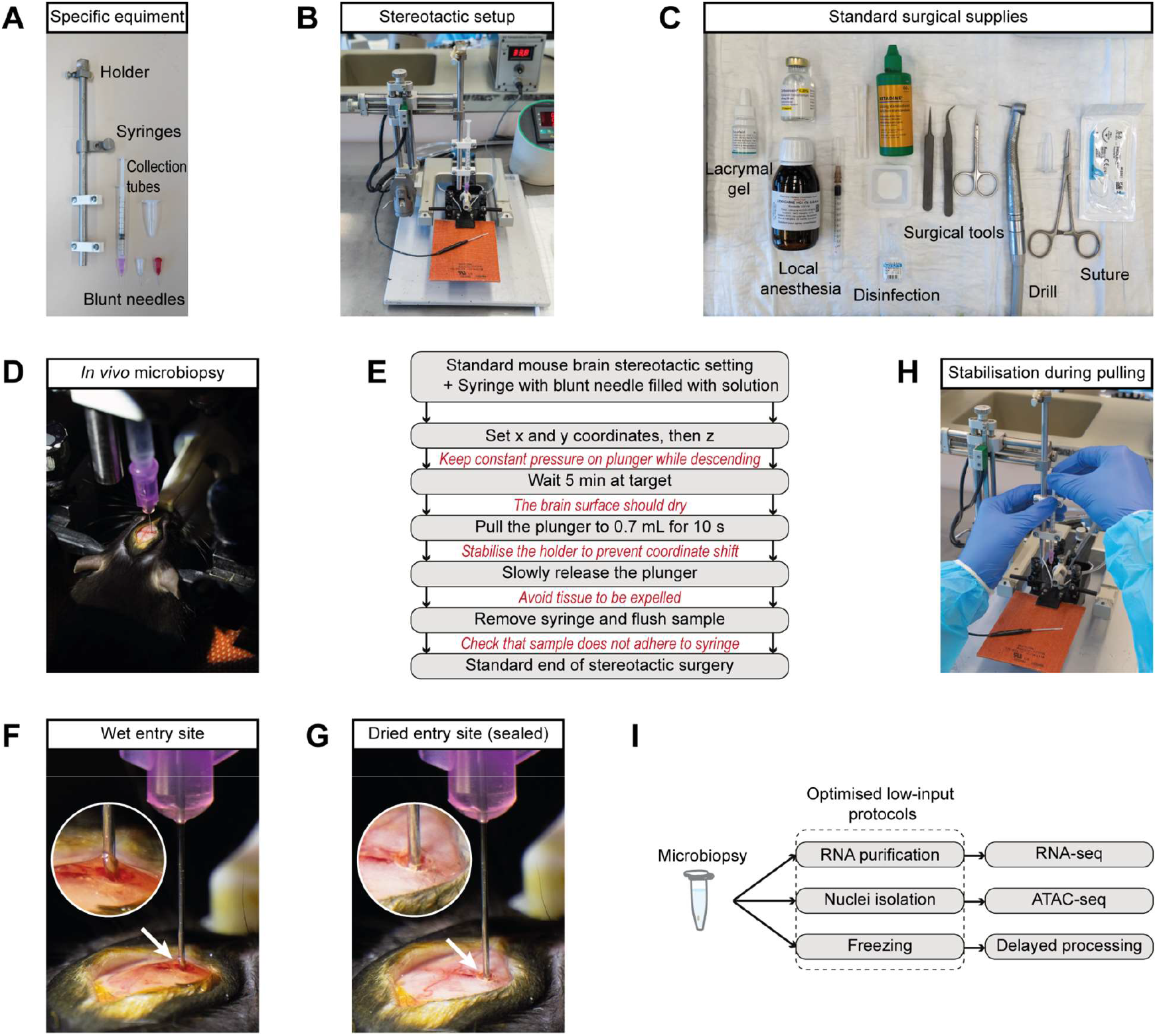
Stereotactic microbiopsy workflow and critical parameters. (A) Specific equipment for microbiopsy. (B) Standard stereotactic frame equipped with the holder, syringe and needle for microbiopsy. (C) Standard surgical supplies used for the procedure. (D) *In vivo* microbiopsy setup before aspiration. (E) Schematic overview of the microbiopsy procedure highlighting critical operational parameters in red. (F) Representative view of the cortical surface before sealing (wet entry site). (G) Sealed, dry entry site that supports local vacuum formation during aspiration. (H) Hand position used to stabilise the syringe during plunger pulling. (I) Schematic of downstream applications.

Immediately after aspiration, the needle/syringe is flushed to transfer the aspirate into a microtube containing the appropriate medium (Fig. 1I). Depending on the intended downstream application, the sample is collected directly into RNA lysis buffer, nuclei isolation buffer for chromatin accessibility profiling, or a freezing medium for later nuclei isolation. Repeated gentle flushing is used to ensure complete recovery, as tissue fragments can adhere to the syringe wall; with 25 G and 27 G needles the aspirate is typically visible during transfer, but not with 30G needles.

### Characterisation and recovery of the biopsy site

We next assessed the extent of the tissue removed by microbiopsy and the recovery. Histological examination of formadelhyde-fixed sections immediately after sampling revealed a spatially restricted biopsy cavity confined to the targeted brain region (Fig. 2A, B). Sampling was performed in a cortical structure (orbitofrontal cortex) and a subcortical structure (dorsomedial striatum), two relatively large regions expected to tolerate limited tissue removal. Higher-magnification images illustrate the aspirated area, with DAPI staining enabling measurement of the cavity diameter.

**Figure 2:**
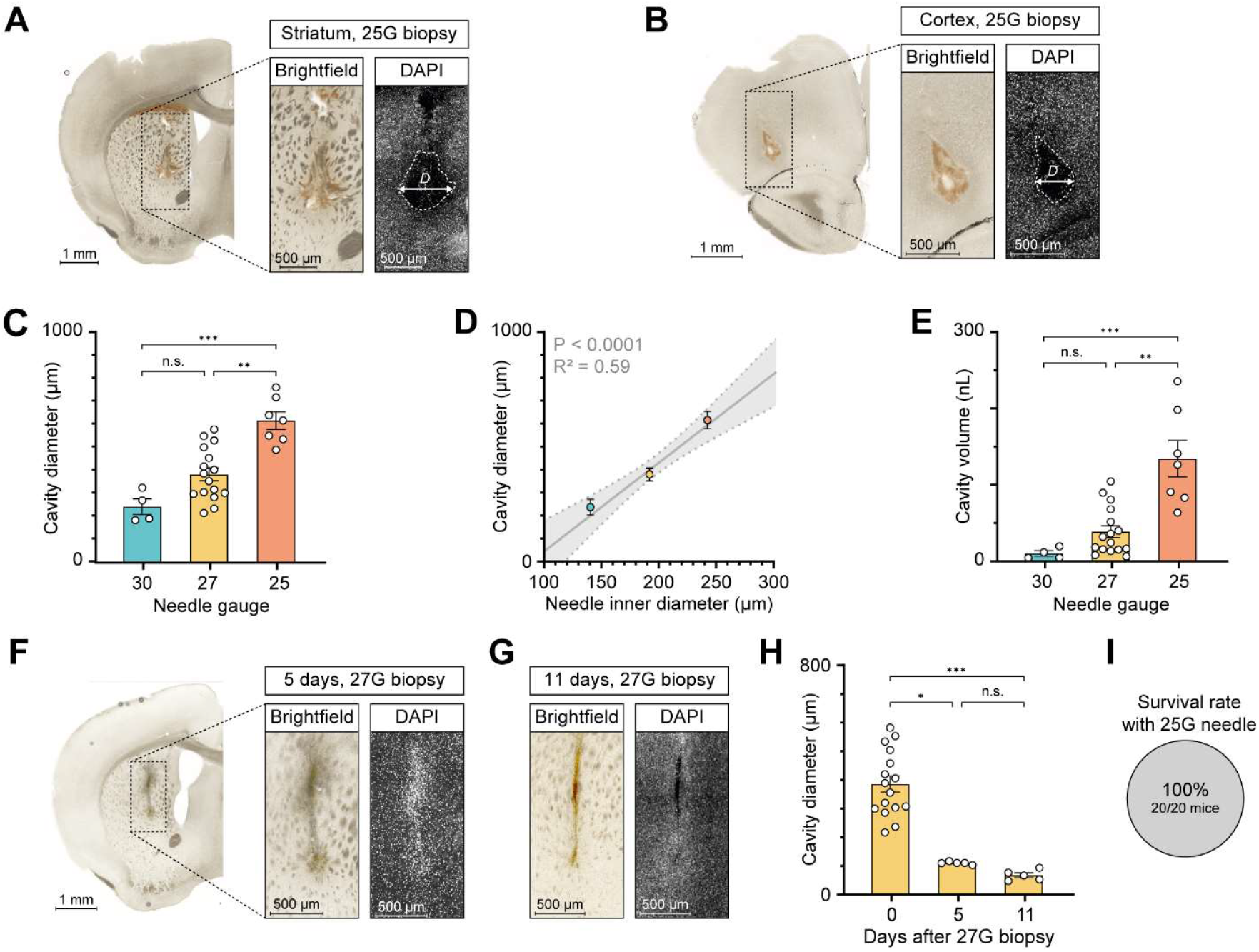
Extent and recovery of the microbiopsy site. (A) Representative histology of a 25G biopsy in the dorsal striatum; D, cavity diameter. (B) Representative histology of a 25G biopsy in the orbitofrontal cortex. (C) Cavity diameter for 30G, 27G, and 25G needles (n = 4, 16, 7) . Kruskal–Wallis test: H(2) = 15.1, P = 0.0005; Dunn’s multiple-comparisons test. (D) Linear regression between needle inner diameter and cavity diameter (n = 4, 16, 7). (E) Estimated cavity volume (assuming a spherical shape) for 30G, 27G, and 25G needles (n = 4, 16, 7). Kruskal–Wallis test: H(2) = 15.1, P = 0.0005; Dunn’s multiple-comparisons test. (F) Representative histology of a 27G biopsy in the dorsal striatum after 5 of recovery. (G) Representative histology of a 27G biopsy in the dorsal striatum after 11 days of recovery. (H) Cavity resorption over time (0, 5, and 11 days) using 27G needles (n = 16, 5, 5). Kruskal–Wallis test: H(2) = 18.9, P < 0.0001; Dunn’s multiple-comparisons test. (I) Survival rate 1 month after 25G unilateral microbiopsy of the orbitofrontal cortex. Data are mean ± SEM. n.s., not significant; *P < 0.05, **P < 0.01, ***P < 0.001. Colour key: 30G, blue; 27G, yellow; 25G, red.

To determine how needle size influences tissue removal, we compared three gauges (25 G, 27 G, and 30 G). Average diameters (mean ± SEM) were 620 ± 37 µm for 25 G, 386 ± 29 µm for 27 G, and 245 ± 34 µm for 30 G (Fig. 2C). The diameter of the aspirated area increased proportionally with needle inner diameter (R^2^ = 0.59, P < 0.0001, Fig. 2D). The corresponding estimated tissue volumes (approximated by a spherical shape) were 133 ± 24 nL, 38 ± 8 nL and 9.0 ± 3.5 nL (Fig. 2E).

We then examined how the biopsy site recovered over time. Separate groups of animals were perfused 5 (Fig. 2F) and 11 days after sampling (Fig. 2G). At 5 days, the biopsy tract contained a compact, DAPI-dense cellular accumulation at the sampling site, consistent with a transient injury response that was no longer evident at day 11. As shown in Fig. 2H, the cavity size decreased markedly over time. Quantification confirmed a 71 % reduction in cavity diameter after 5 days and 82% after 11 days relative to the immediate post-biopsy condition, i.e. an estimated reduction of cavity volume of 98 and 99%.

Animals monitored postoperatively recovered normally with no procedure-related mortality. This included a separate cohort of 20 mice that underwent 25G unilateral orbitofrontal cortex microbiopsy and were maintained for >1 month before sacrifice (Fig. 2I). Mice exhibited no abnormal behaviour or signs of discomfort according to postoperative welfare scoring, comparable to animals undergoing standard stereotactic procedures. These findings demonstrate that stereotactic microbiopsy removes a well-confined, reproducible tissue volume that undergoes rapid structural recovery, allowing subsequent behavioural or molecular analyses in the same animal.

### Isolation and quantification of nuclei from microbiopsy samples

We next evaluated whether microbiopsies yield sufficient nuclei for downstream molecular assays. Microbiopsies were processed either immediately or after cryopreservation in an optimised sampling/freezing solution followed by thawing. Tissue fragments were mechanically dissociated using a Dounce homogenizer and nuclei were quantified. Fluorescence microscopy confirmed intact DAPI-positive nuclei after dissociation of frozen–thawed microbiopsies (Fig. 3A). Cryopreservation did not measurably reduce nuclei recovery, with comparable yields from fresh and frozen samples (Fig. 3B). Nuclei yields increased with needle gauge, ranging from 540 ± 70 (mean ± SEM) nuclei with 30 G needles to 1,200 ± 150 with 27 G needles and 3,100 ± 800 with 25 G needles (Fig. 3C). Consistent with this, nuclei recovery scaled with the estimated cavity volume (R^2^ = 0.53, P < 0.001; Fig. 3D), supporting a predictable relationship between aspirated tissue volume and nuclei yield.

**Figure 3:**
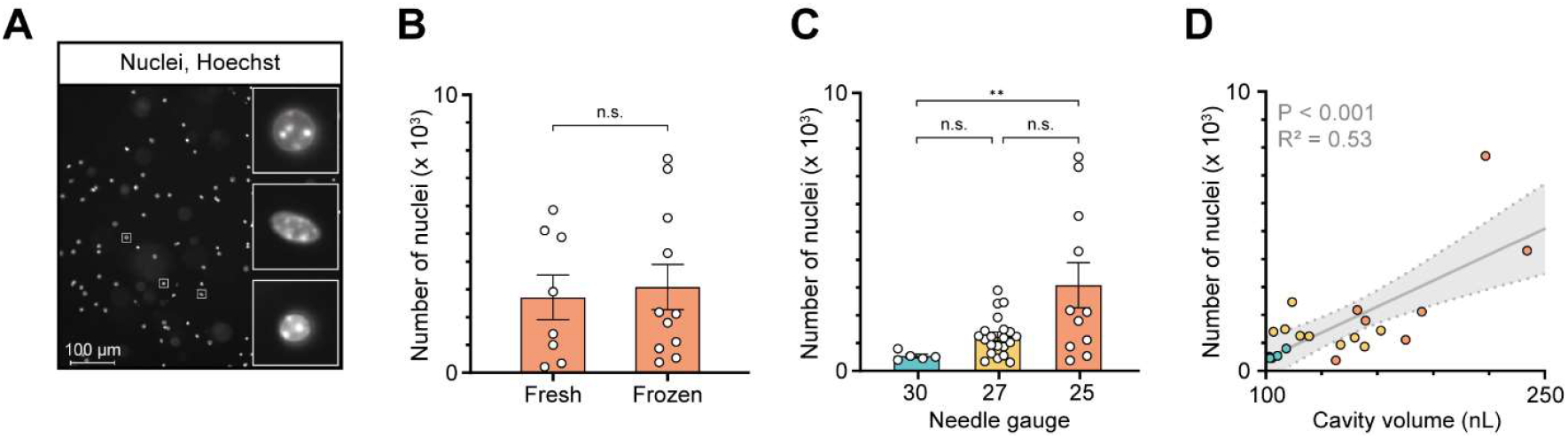
Freezing and quantifying nuclei. (A) Representative image of nuclei isolated from a microbiopsy after freezing, thawing, and dissociation; magnified insets illustrate preserved nuclear integrity. (B) Quantification of nuclei from fresh versus frozen 25G microbiopsies (n = 8, 11). Mann–Whitney U test (one-tailed): U = 39, P = 0.36. (C) Quantification of nuclei obtained from frozen microbiopsies using 30G, 27G, and 25G needles (n = 5, 21, 11). Kruskal–Wallis test: H(2) = 8.8, P = 0.012; Dunn’s multiple-comparisons test. (D) Linear regression between estimated cavity volume (calculated from cavity diameter) and number of nuclei (n = 21). Data are mean ± SEM; n.s., not significant; **P < 0.01. Colour key: 30G, blue; 27G, yellow; 25G, red.

### RNA yield and chromatin integrity from microbiopsy samples

To enable transcriptomic and epigenomic assays, we established two downstream applications from the microbiopsy samples: RNA extraction and nuclei purification followed by ATAC-seq. For RNA extraction, 1 mL of RLT buffer supplemented with 1 % β-mercaptoethanol was added directly to each sample. After brief vortexing and addition of 700 μL of ethanol 100%, RNA was purified on column with DNase digestion using the Qiagen RNeasy Micro Kit. RNA quantity and integrity were assessed from single microbiopsies collected with 27 G needles. Total RNA yield per sample ranged from 1–13 ng (Fig. 4A). RNA integrity numbers (RIN) obtained using a Bioanalyzer (Agilent) system were consistently above 7 (Fig. 4B), and electropherograms displayed clear 18S and 28S peaks (Fig. 4C), indicating minimal degradation. These yields and quality metrics fall within the range suitable for bulk RNA-seq library preparation and sequencing.

**Figure 4:**
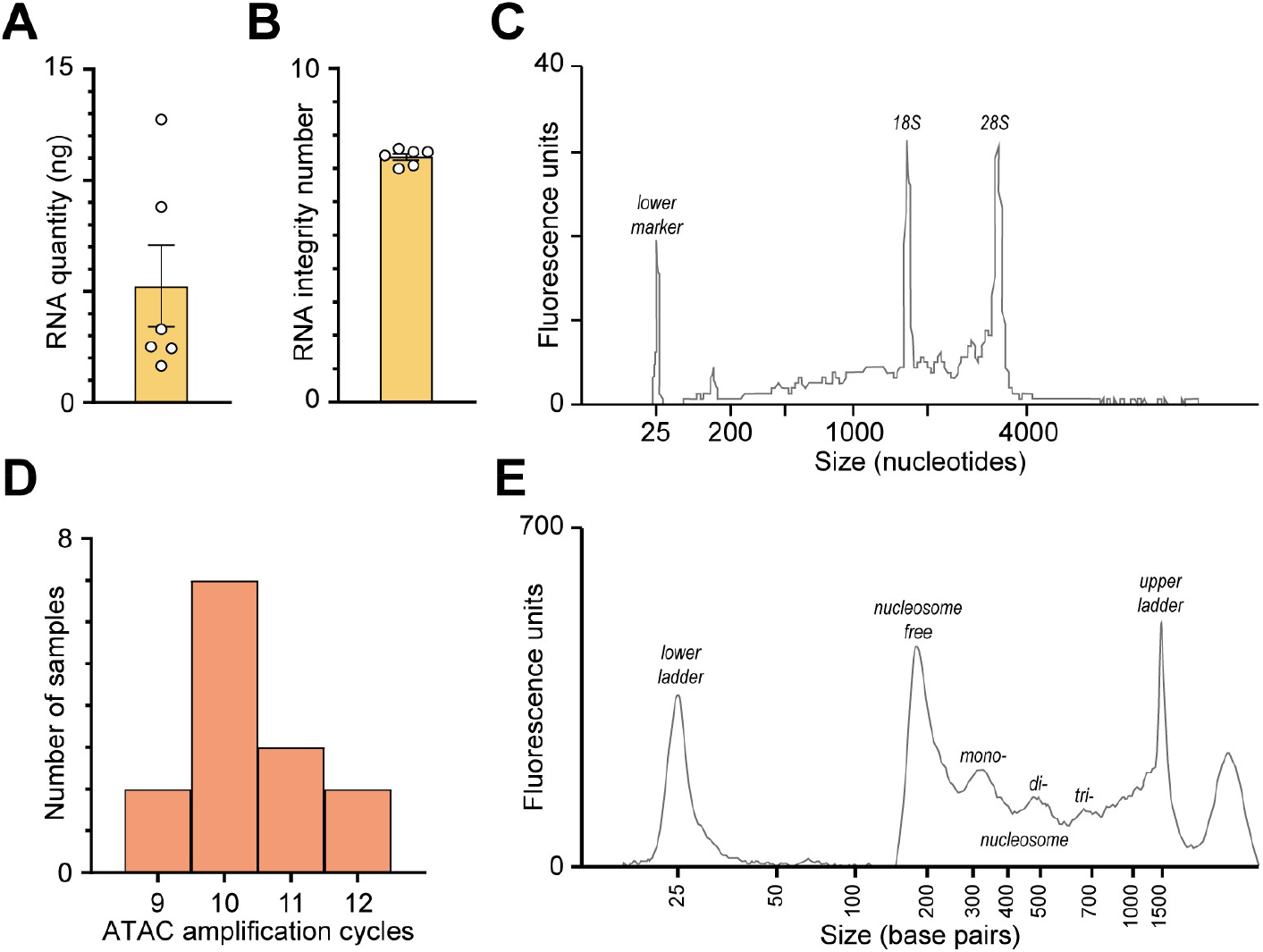
Downstream applications: RNA and chromatin accessibility. (A) RNA yield from purified total RNA isolated from 27G microbiopsies (n = 6). (B) RNA integrity of RNA isolated from 27G microbiopsies (n = 6). (C) Representative RNA electropherogram (Bioanalyzer) illustrating RNA quality. (D) Distribution of the total number of PCR amplification cycles required for ATAC library preparation from 25G microbiopsies (n = 14). (E) Representative ATAC-seq library DNA electropherogram (TapeStation) illustrating fragment-size distribution. Data are mean ± SEM.

For chromatin accessibility analysis, we developed an in-house nuclei purification and ATAC-seq workflow optimised for small input volumes. The purification protocol, adapted from Marion-Poll et al. ^16^, included Dounce homogenisation, permeabilisation with igepal, and purification by centrifugation through an iodixanol density solution. Transposition and library preparation were performed following Buenrostro *et al*. ^4^, with total PCR amplification cycles ranging from 9–12 (Fig. 4D), within standard parameters. Fragment-size distributions (TapeStation, Agilent) showed the expected nucleosomal periodicity (Fig. 4E), confirming preservation of chromatin structure and accessibility.

Together, these analyses demonstrate that microbiopsy samples yield high-quality RNA and nuclei suitable for both RNA-seq and ATAC-seq from nanoliter-scale volumes of mouse brain tissue.

### Technical parameters influencing sampling accuracy and efficiency

During method development, we identified two parameters that were critical for sampling purity and targeting precision. First, maintaining positive pressure while advancing the needle towards the target prevented tissue located above the target from entering the lumen prematurely. In the absence of counterpressure, off-target tissue could enter the needle tract, resulting in contamination (Fig. 5A). In contrast, when counterpressure was maintained, no contamination was observed (Fig. 5B).

**Figure 5:**
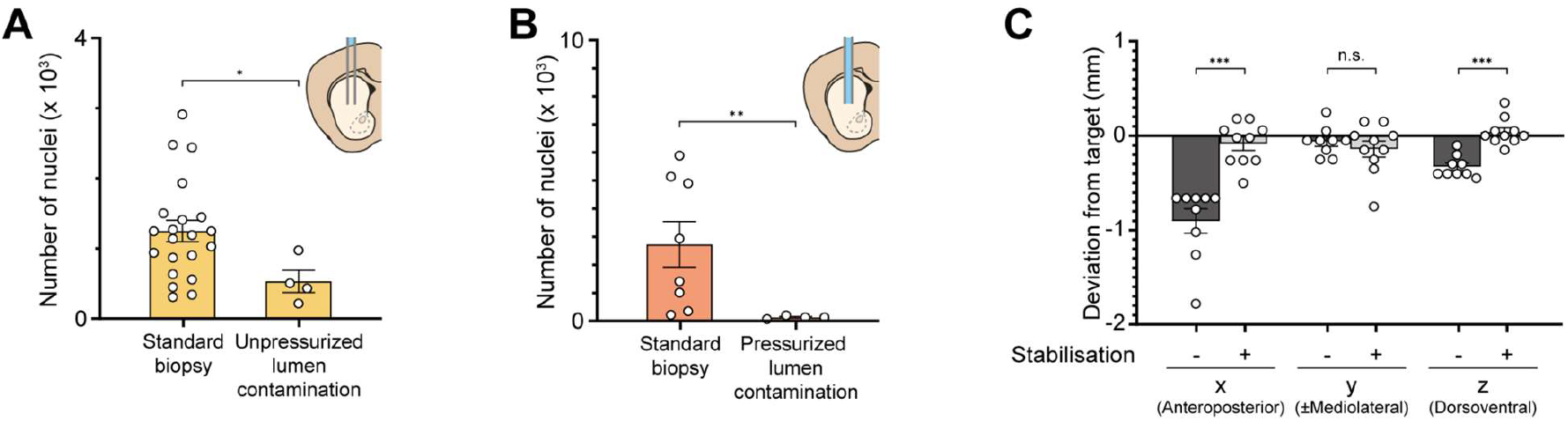
Technical parameters influencing sampling purity and targeting precision. (A) Nuclei yield from standard 27G frozen microbiopsies versus nuclei recovered from tissue that entered the unpressurized lumen in the absence of aspiration (n = 21, 4) . Mann–Whitney U test (one-tailed): U = 13, P = 0.015. (B) Nuclei yield from standard 25G fresh microbiopsies versus a needle insertion performed under positive pressure without subsequent aspiration (n = 8, 4). Mann–Whitney U test (one-tailed): U = 0, P = 0.002. (C) Absolute deviation from the intended target along the stereotactic axes, comparing biopsies performed with versus without stabilisation (n = 9,10). Mann–Whitney tests with FDR correction. Data are mean ± SEM; n.s., not significant; *P < 0.05, **P < 0.01, ***P < 0.001.

Second, the mechanical force generated during aspiration could slightly displace the needle tip and shift the effective stereotaxic coordinates. Stabilising the syringe with the contralateral hand (Fig. 1H) to counteract this force markedly improved targeting accuracy (Fig. 5C). The mean anteroposterior deviation was −0.90 mm without stabilisation and −0.08 mm with stabilisation.

Together, these observations define key operational parameters for reliable microbiopsy performance: continuous low positive pressure during descent and manual stabilisation of the syringe during aspiration, in addition to the other procedural recommendations summarised in Fig. 1C. Adherence to these steps improves sampling specificity, preserves targeting accuracy, and maximises recovery of high-quality material.

## DISCUSSION

This study introduces a stereotactic microbiopsy technique that enables recovery of intracellular material from spatially defined regions of the living mouse brain while preserving postoperative survival. By combining standard stereotactic surgery with nanoliter-scale aspiration, the method allows isolation of RNA or intact nuclei suitable for bulk RNA-seq and ATAC-seq. Importantly, microbiopsy enables molecular sampling prior to experimental manipulation, supporting future within-subject longitudinal designs that are otherwise not feasible in mice.

A principal methodological advance will be the ability to dissociate baseline molecular state from experience- or treatment-induced change. Most neuroscience molecular studies in rodents rely on cross-sectional comparisons, in which post-intervention differences may reflect either preexisting variability or induced effects. By permitting pre-intervention sampling from the same individual, microbiopsy controls between-animal variance and increases statistical power for detecting biologically meaningful changes. This design is particularly relevant in contexts where inter-individual molecular heterogeneity is substantial, even among genetically identical animals, and where baseline state may shape subsequent behavioural, physiological, or pathological outcomes.

Microbiopsy addresses a gap between existing *in vivo* sampling approaches. Clinical and large-animal stereotactic biopsies provide access to intracellular material but remove tissue volumes that are incompatible with the mouse brain. Conversely, rodent techniques such as microdialysis^10^, solid-phase microextraction^11^, and open-flow microperfusion^12^ enable repeated sampling but are restricted to extracellular analytes and do not recover nucleic acids. The approach described here provides a minimally invasive route to retrieve intracellular RNA and chromatin from defined mouse brain regions, using widely available stereotactic equipment and straightforward sample handling. It should be readily adaptable to rats, which have larger brains and are routinely used for stereotactic neurosurgical studies.

Reliable performance depended on a defined set of operational parameters. Maintaining positive pressure during needle descent prevented premature entry of tissue into the lumen and reduced off-target contamination. A 5-minute pause at the target coordinate enabled local sealing of the entry site and consistent vacuum formation during aspiration. Mechanical stabilisation of the syringe during plunger retraction minimised needle deflection and substantially improved targeting accuracy. After aspiration, slow release of the plunger without forward pressure and repeated gentle flushing were required to ensure complete recovery of the tissue fragment and to avoid loss through adhesion to the syringe wall. Together, these parameters define practical controls that materially affect sampling purity, yield, and anatomical precision.

A key biological consideration is the influence of anaesthesia. Isoflurane and related agents are known to alter transcriptional and epigenomic states^17-19^, and may therefore contribute to differences between microbiopsy-derived profiles and profiles obtained from terminal collections. For longitudinal designs, this favours standardising anaesthetic exposure across time points or interpreting the procedure itself as a controlled perturbation. In this context, microbiopsy is best viewed as enabling relative comparisons within individuals rather than absolute equivalence to terminal sampling.

The method is likely to be most appropriate for relatively large brain regions that can tolerate limited tissue removal. Although histological analysis indicated recovery over time, tissue removal inevitably induces a local perturbation, and resilience may not generalise to small or highly specialised nuclei. While the approach could in principle support repeated sampling, this was not evaluated here due to ethical considerations and the risk of cumulative damage or gliosis at the same site.

Within neuroscience, stereotactic microbiopsy should be applicable to a wide range of experimental contexts in which intra-individual molecular trajectories are of interest, including pharmacological interventions, disease models with variable penetrance or progression, and ageing paradigms.

In addition to enabling within-subject molecular designs, microbiopsy has ethical implications. Longitudinal sampling reduces reliance on terminal endpoints and could decrease the number of animals required for multi-timepoint molecular studies, consistent with principles of refinement and reduction.

In summary, stereotactic microbiopsy provides a reproducible and minimally invasive strategy to obtain RNA and chromatin information from the living mouse brain. By enabling longitudinal molecular sampling within individuals, the method supports experimental designs that distinguish baseline molecular state from induced change, while remaining compatible with standard stereotactic workflows and low-input sequencing technologies.

## METHODS

### EXPERIMENTAL MODEL

Experiments were performed on adult female and male C57BL/6J mice (Charles River Laboratories). Animals were housed in groups under standard conditions (12-h light/dark cycle, controlled temperature) with food and water available *ad libitum*. All procedures were conducted in accordance with institutional and national ethical guidelines and were approved by the Institutional Animal Care and Use Committee of the University of Geneva and by the animal welfare committee of the Canton of Geneva (protocol numbers GE336B and GE362A).

### METHOD DETAILS

#### Sampling and freezing solution

The solution used for sampling and freezing consisted of 27% (w/v) sucrose prepared in artificial cerebrospinal fluid (ACSF) composed of 119 mM NaCl, 2.5 mM KCl, 1.3 mM MgCl_2_, 2.5 mM CaCl_2_, 11 mM glucose, 26.2 mM NaHCO_3_, 1 mM NaH_2_PO4, and 789 mM sucrose, using RNase-free reagents. The solution was oxygenated for 30 min with 95% O_2_ / 5% CO_2_, and the pH was verified to be within 7.3–7.4. It was then sterile-filtered through a 0.22 µm membrane (Stericup, Millipore). Aliquots of 300 µL were prepared in RNase-free 1.5 mL tubes (Eppendorf Clearlock), and heat-sterilised at 95 °C for 5 min using a ThermoMixer (Eppendorf). After a brief centrifugation, aliquots were stored at –20 °C until use.

This sucrose-based ACSF is sterile, physiologically compatible with brain tissue, and free of RNA-degrading agents. Its high sucrose concentration provides cryoprotection, allowing aspirated material or isolated nuclei to be frozen directly without additional processing.

#### Stereotactic microbiopsies

Mice were anesthetised with 4% isoflurane in O_2_ for induction and maintained at 2% isoflurane via a nose cone throughout the procedure. The absence of blink, toe-pinch, and tail reflexes was verified before fixation in a stereotactic frame (Stoeling). A mixture of lidocaine 0.5% and bupivacaine 0.25% (120–160 µL) was applied subcutaneously at the incision site for local anesthesia. Animals were placed on a heating pad to maintain body temperature at 37 °C, and lacrimal gel (Lacryvisc, Alcon, Switzerland) was applied to prevent corneal dehydration.

The scalp was disinfected with 10% povidone-iodine solution (Betadine) before incision, and aseptic surgical instruments were used throughout. After a midline incision, the exposed skull surface was disinfected again, and the landmarks bregma and lambda were identified. A 1 mm craniotomy was then made at the target location using a surgical drill. Target coordinates were referenced to bregma and measured from the cortical surface: +0.8 mm AP, 1.65 mm ML, 2.9 mm DV for the dorsomedial striatum, and +2.6 mm AP, 1.75 mm ML, 1.55 mm DV for the orbitofrontal cortex. The dorsoventral coordinate for each site corresponds to the target depth minus 400 µm, compensating for the upward draw of tissue that occurs during aspiration.

Microbiopsies were performed using sterile blunt-tipped needles (25 G, 27 G, or 30 G, 0.5 inch long; Metcal) attached to 1 mL syringes prefilled with the sampling solution. All air bubbles were expelled beforehand to maintain continuous fluid contact, ensure stable vacuum formation, and prevent inadvertent air injection that could damage tissue.

During needle descent, a positive pressure was applied on the plunger to prevent brain tissue from entering the needle lumen prematurely. At the target coordinate, pressure on the syringe plunger was released to allow tissue adhesion and sealing. The needle was left in place for 5 min to enable localised sealing. If a larger volume of solution had been expelled during descent, the waiting time was extended until the cortical surface appeared dry and sealed against the needle.

Suction was generated by rapidly retracting the syringe plunger to 0.7–0.9 mL, holding this position for 10 seconds, and then slowly releasing it over ∼5 seconds while restraining the plunger to prevent a sudden forward return that could create positive pressure. Forward pressure was avoided to prevent expelling the aspirated tissue. Because the applied force can cause needle deflection, the syringe was stabilised with the opposite hand during aspiration to maintain targeting accuracy. Failure of the plunger to return close to its starting position upon release was taken as an indication of incomplete sealing between the entry tract and the cortical surface.

The syringe and needle were then withdrawn and detached from the holder. For RNA extraction, the aspirated tissue was expelled by three gentle flushes of the syringe into a 2 mL RNase-free microcentrifuge tube prefilled with 1mL RLT lysis buffer (RNeasy Micro Kit, Qiagen) supplemented with 1% β-mercaptoethanol. For nuclei freezing, the sample was flushed three times into an empty 1.5 mL tube, while for ATAC-seq applications, the tissue was expelled into a 1.5 mL tube containing 700 µL homogenisation buffer on ice (see corresponding subsection). To recover residual material, a small volume of air was aspirated into the syringe at the end of the procedure; the needle was then removed, and the remaining contents of the syringe were expelled into the collection tube.

The scalp incision was closed with absorbable sutures (Prolene 5-0). For animals not perfused immediately after sampling, postoperative hydration was monitored using the skin-pinch test, and mice showing signs of dehydration received 0.5 mL sterile 0.9% NaCl subcutaneously. Postoperative analgesia was provided by carprofen (Rimadyl) in the drinking water (0.05 mg/mL; 0.1% v/v) for two days, and animals were monitored daily for three days following surgery.

#### Freezing microbiopsies

When samples were not processed immediately, tubes containing microbiopsies in sampling/freezing solution or RLT buffer (for RNA extraction) were placed on dry ice for rapid freezing and subsequently stored at –20 °C for nuclei analysis or at –80 °C for RNA extraction. Prior to further processing, samples were thawed on ice.

#### RNA extraction

RNA was extracted using the RNeasy Micro Kit (Qiagen) with minor adjustments to accommodate microbiopsy samples. Each biopsy (approximately 300 µL of sampling buffer containing the tissue) was mixed with 1 mL of Buffer RLT supplemented with 1% β-mercaptoethanol, vortexed to lyse the tissue, and briefly centrifuged. Seven hundred microliters of 100% ethanol were then added (final ethanol concentration ≈35%), mixed by vortexing, and briefly spun down.

The lysate, including any precipitate, was applied to a RNeasy Micro spin column in ≈700 µL aliquots and centrifuged for 15 seconds at ≥8,000 g after each load. The flow-through was discarded between steps until the full sample volume had passed through the column. Washing steps followed the manufacturer’s instructions, beginning with Buffer RW1, and included an on-column DNase digestion to remove genomic DNA. After the final Buffer RPE wash, columns were centrifuged for an additional 2 minutes to ensure complete removal of residual wash solution.

RNA was eluted in 12.5 µL of RNase-free water and immediately placed on ice. RNA concentration and integrity were assessed using an Agilent Bioanalyzer RNA Pico Kit, and both RNA integrity numbers (RIN) and concentrations were measured for each sample.

#### Transposition of nuclei

All buffers were prepared from DNase-free reagents and filtered through a 0.22 µm membrane (Stericup, Millipore) before use, except the transposition mix. The homogenisation buffer contained 250 mM sucrose, 25 mM KCl, 5 mM MgCl_2_, 20 mM Tris-HCl (pH 7.4), 1% (w/v) BSA, and 0.1% (v/v) Tween-20. The working buffer consisted of 150 mM KCl, 30 mM MgCl_2_, and 120 mM Tris-HCl (pH 7.4). OptiPrep (60% iodixanol solution) was mixed with the working buffer (5:1, v/v) to prepare a 50% iodixanol solution, which was then diluted 1:1 with homogenisation buffer to obtain a 25% iodixanol solution. A wash buffer composed of 20 mM Tris-HCl (pH 7.4), 10 mM MgCl_2_, and 1% BSA was used for the final rinse. The transposition mix was freshly prepared for each sample (total 50 µL) by combining 25 µL TD buffer 2× (Illumina), 22.5 µL nuclease-free water, and 2.5 µL Tn5 transposase (Illumina). Low binding DNase free tips were used throughout the protocol.

The purification procedure was adapted from Marion-Poll *et al*. ^16^ to minimise handling and preserve nuclei integrity in small, myelin-rich brain samples. Approximately 300 µL of sampling buffer containing the tissue was added to 700 µL of homogenization buffer on ice and immediately transferred to a 2 mL Dounce homogenizer, without pre-mixing. Homogenization was performed with 12-20 strokes of pestle B (clearance 12–63 µm) to gently dissociate tissue and release intact nuclei. The homogenate was centrifuged at 2,000 × g for 5 min at 4 °C, and the pellet was gently resuspended in homogenisation buffer containing 0.1% Igepal and a 5 min timer was started. Resuspension was performed by pipetting up and down 15 times, carefully aspirating material from the bottom of the tube and expelling it against the wall while keeping the pipette tilted, without touching the tube and avoiding bubble formation. This technique should always be used when resuspending nuclei to preserve integrity and minimise shear damage.

A 300 µL layer of 25% iodixanol was then placed beneath the suspension using a syringe and needle. At the end of the 5 minutes igepal incubation, samples were centrifuged at 10,000 × g for 30 min at 4 °C, the supernatant was discarded, and 300 µL of fresh 25% iodixanol was added without resuspending the pellet. A second centrifugation was performed at 10,000 × g for 5 min at 4 °C. The pellet was then washed with 1 mL of wash buffer, centrifuged at 2,000 × g for 5 min at 4 °C, and the supernatant was removed.

The purified nuclei were resuspended in 50 µL of transposition mix by gentle pipetting 15 times. Transposition was performed for 30 min at 37 °C in a thermomixer without agitation. Samples were then frozen at –20 °C until further processing.

Subsequent steps followed the Buenrostro *et al*. ^4^ ATAC-seq protocol. DNA was purified using the Qiagen MinElute Kit, pre-amplified for five PCR cycles (including an initial 5 min extension at 72 °C before denaturation), and the number of additional cycles (typically 4–7, total 9–12) was determined by qPCR. Final libraries were purified with MinElute columns, and fragment size distributions were verified on an Agilent Tapestation, confirming the expected nucleosomal periodicity before potential sequencing.

#### Perfusion and fixation of mice

Animals were deeply anesthetised with 150 mg/kg pentobarbital (Esconarkon, 10 mL/kg i.p.) and transcardially perfused with 4% (wt/vol) formaldehyde in PBS (pH 7.4) at 20 mL/min at room temperature for 5 min. Brains were post-fixed overnight at 4°C before being cut in 80 μm coronal sections in PBS using a vibratome (Leica, VT1200S). The slides were mounted in Vectashield with DAPI (Vector Laboratories, USA).

#### Imaging brain sections and estimating cavity size

Images were acquired using an automated slide scanner with a 10X objective (Zeiss Axioscan Z1), in brightfield and DAPI fluorescence. Cavity diameter was measured as the maximal diameter of the cavity in the section showing the largest cavity area, using the ZEISS ZEN lite software.

#### Localisation of biopsy site and comparison to target point

For localisation of the biopsy site we identified the center of the cavity. The coronal sections were aligned with the mouse brain atlas^15^ and position measured. Targeting error was calculated as measured coordinates minus intended target coordinates for each axis.

#### Nuclei quantification

Microbiopsies were processed either immediately or after cryopreservation. Frozen samples were thawed on ice. Samples (∼300 µL of sampling buffer containing the tissue) were brought to 1 mL by adding PBS containing 1% (w/v) BSA.. The sample was then mechanically dissociated using a 2 mL Dounce homogeniser, applying 12-20 strokes of pestle B (clearance 12–63 µm). The homogenate was centrifuged at 2,000 × g for 5 min at 4 °C, and the pellet was gently resuspended in PBS–BSA (1%) containing Hoechst 33258 0.05 µg/mL.

Aliquots were transferred into black, optical-bottom 96-well microplates (Greiner Bio-One). Imaging was performed using an ImageXpress automated microscope (Molecular Devices). Nuclei counting was automated using the MetaXpress (Version 6.7.2.290), applying standard nuclei count with a lower threshold of 5 µm and an upper threshold of 15 µm in diameter.

### QUANTIFICATION AND STATISTICAL ANALYSIS

All statistical analyses were performed using GraphPad Prism (version 10). Details of statistical tests are provided in the corresponding figure legends. Data are presented as mean ± SEM. Individual data points represent independent microbiopsies. No data points were excluded from analysis.

For comparisons between two groups, nonparametric Mann–Whitney tests were used. One-tailed tests were applied when a directional effect was defined a priori based on experimental design. For comparisons among more than two groups, Kruskal–Wallis tests were performed followed by Dunn’s multiple-comparisons tests with adjusted *P* values. For the analysis of deviations along stereotaxic axes, which involved multiple related comparisons, false discovery rate (FDR) correction was applied using the Benjamini–Krieger–Yekutieli procedure with a desired FDR (Q) of 1%.

Linear regression analyses were performed using least-squares fitting, and significance of the slope was assessed by F test. Goodness of fit is reported as R^2^.

Statistical significance was defined as *P* < 0.05. Significance levels are indicated in the figures as follows: ns, not significant; \**P* < 0.05; \*\**P* < 0.01; \*\*\**P* < 0.001.

## RESOURCE AVAILABILITY

### Lead contact

Requests for further information and resources should be directed to and will be fulfilled by the lead contact, Lucile Marion-Poll.

### Materials availability

This study did not generate new unique reagents.

### Data and code availability

- Source Data is provided as an excel file.
- All data reported in this paper will be shared by the lead contact upon request.
- Any additional information required to reanalyse the data reported in this paper is available from the lead contact upon request.
- This paper does not report original code.

## ACKNOWLEDGMENTS

We thank Jennifer Cand for the animal care, and members of the Lüscher lab for stimulating discussions. Microscopy acquisition and analysis were performed at the Bioimaging Core Facility, Faculty of Medicine, University of Geneva. Electrophoresis profiles were generated by the iGE3 Genomics Platform of the University of Geneva. This work was supported by Carigest SA (anonymous donor). Alexander Hoyningen was funded by a MD-PhD fellowship from the Swiss National Science Foundation (SNSF, 323630_214535).

## AUTHOR CONTRIBUTIONS

Conceptualisation, L.M.-P., and C.L..; Methodology, A.Hoyningen. and L.M.-P.; Investigation, A.Hoyningen, L.M.-P., L.F., A.Hiver; Visualisation, A.Hoyningen, A.L. and L.M.-P.; Validation A.R. and L.M.-P.; Writing— original draft, A.Hoyningen and L.M.-P.; Writing—review & editing, A.L., A.R. and C.L.; Funding acquisition, L.M.-P. and C.L.; Resources, C.L.; Supervision, L.M.-P. and C.L.

## DECLARATION OF GENERATIVE AI AND AI-ASSISTED TECHNOLOGIES

During the preparation of this work, the author(s) used ChatGPT 5.2 for language editing. After using this tool, the author(s) reviewed and edited the content as needed and take full responsibility for the content of the publication.

